# Distinct neural processing of real versus unreal optic flow in the human brain

**DOI:** 10.64898/2025.12.31.696655

**Authors:** Xuechun Shen, Zhoukuidong Shan, Shuguang Kuai, Li Li

## Abstract

Visual perception of self-movement relies on optic flow, a global motion pattern generated by self-movement in a rigid 3D environment. Such optic flow conveys both 2D features (e.g., radial velocity fields and centers of motion) and 3D structural information (e.g., motion perspective). However, many previous neuroimaging studies have used simplified 2D radial motion as “real” optic flow, leaving it unclear whether such “unreal” stimuli adequately capture how the brain processes ecologically valid self-movement information. Previous behavioral work has shown that sensitivity to expansion and contraction reverses depending on whether optic flow is ecologically valid (L. Li et al., 2025), raising the question of how such differences are represented in the human brain.

We conducted two functional magnetic resonance imaging (fMRI) experiments to systematically compare cortical processing of real versus unreal optic flow. The two stimulus types were strictly matched in 2D global features and local motion signals, differing only in whether they preserved 3D structure consistent with self-movement. To minimize task-related influences, participants performed a task-irrelevant fixation point color-change detection task during scanning. Experiment 1 used a block design in which participants viewed expanding and contracting optic flow at different motion coherence levels. Experiment 2 presented real and unreal optic flow within the same scanning session to directly test neural sensitivity to ecological validity.

Multivoxel pattern analysis (MVPA) revealed that, among multiple visual and optic flow-responsive regions, the human dorsal medial superior temporal area (MST) exhibited robust and consistent sensitivity to the ecological validity of optic flow. Specifically, under real optic flow, decoding accuracy for motion coherence in MST was significantly higher for expansion than for contraction, whereas this preference was reversed under unreal optic flow. Neural threshold estimates derived from fMR-metric functions further supported this pattern reversal, indicating systematic modulation of MST sensitivity by ecological structure. Experiment 2 further demonstrated that, regardless of motion pattern, MST reliably distinguished real from unreal optic flow, with decoding accuracy increasing as motion coherence increased. These findings indicate that MST encodes not only motion patterns themselves but also whether optic flow conforms to the ecological structure of self-movement.

Together, these results indicate that MST responses are not determined solely by 2D radial motion features, but are modulated by whether optic flow preserves 3D structure consistent with self-movement. This work clarifies a long-standing conflation between simplified radial motion and real optic flow and highlights the importance of ecological structure in shaping dorsal-stream motion representations related to self-movement perception.

## Introduction

When observers move through the environment, self-movement generates a global pattern of retinal motion known as optic flow (Gibson, 1950). During forward or backward translation, optic flow conveys not only 2D global features, such as a radial velocity field and a center of motion (CoM), but also 3D structural information, including motion parallax and perspective cues. Together, these signals specify the direction (i.e., heading) and speed of self-movement (Gibson, 1950; Matthis et al., 2022; R. Warren, 1976; W. H. Warren, 2006) and provide information about the spatial layout of the environment (Koenderink, 1986; Muller et al., 2023; Rogers, 2021; Todd, 1995), thereby supporting navigation-related behaviors such as obstacle avoidance and postural control.

A large body of work in humans and non-human primates has identified a distributed cortical network involved in optic flow processing, spanning temporal, parietal, and cingulate regions, including MT, MST, V3A, V6, V7, KO/V3B, CSv, precuneus (Pc), and VIP (e.g., Cardin et al., 2012; Cardin & Smith, 2010, 2011; Greenlee, 2000; Koyama et al., 2005; Morrone et al., 2000; Smith et al., 2006; Wall & Smith, 2008). However, many of these studies have relied on simplified “unreal” stimuli that mimic certain 2D motion features (e.g., a radial velocity field and CoM) but do not preserve the 3D structure generated by self-movement in a rigid environment. For example, some studies used radial random-dot motion patterns in which all dots moved toward or away from the CoM at a constant speed (e.g., Cardin et al., 2012; Cardin & Smith, 2010; Koyama et al., 2005; Morrone et al., 2000; Wall & Smith, 2008), or in which dot speed increased linearly with distance from the CoM on the image plane (e.g., Graziano et al., 1994; Greenlee, 2000; Rutschmann et al., 2000; Tanaka & Saito, 1989). Because such stimuli lack key self-movement-consistent structure, it remains unclear whether the neural mechanism reported in these studies fully generalize to natural self-movement perception. This raises a fundamental question: does the visual system process ecologically valid optic flow that preserves the 3D structural regularities generated by self-movement in a rigid environment, in a manner distinct from simplified 2D radial motion?

This question has recently been addressed at the behavioral level. A psychophysical study compared detection sensitivity to expansion and contraction in real versus unreal optic flow while carefully matching 2D global features and local motion signals (Experiment 2 in Li et al., 2025). In that study, participants performed a motion pattern detection task across different coherence levels (signal dot ratios), reporting whether any coherent motion was perceived following each stimulus presentation. The results revealed a clear reversal pattern: under unreal optic flow, participants were more sensitive to contraction than expansion, whereas under real optic flow, sensitivity was higher for expansion than contraction. These findings indicate that the visual system adopts distinct processing strategies depending on whether optic flow is ecologically valid.

Importantly, prior work on heading perception provides indirect neural evidence consistent with this distinction. Disrupting dorsal-stream regions such as MST using transcranial magnetic stimulation (TMS) does not affect heading judgments based on unreal optic flow (Strong et al., 2017), whereas applying TMS to MST during viewing of real ground-plane optic flow significantly increases the variance of heading judgments and reduces their precision (Schmitt et al., 2020). Similarly, single-unit recordings in non-human primates show that neurons in MSTd encode self-movement and support accurate heading estimation under real optic flow conditions (Duffy & Wurtz, 1995; Gu et al., 2012). However, these studies primarily focused on heading estimation and did not directly examine how ecological validity shapes cortical representations of motion patterns when 2D global features and local motion signals are matched. Thus, the neural mechanisms underlying the behavioral dissociation between real and unreal optic flow, particularly the reversal in sensitivity to expansion and contraction, remain unknown.

The present study directly tested whether ecological validity reshapes cortical representations of motion coherence and whether the human brain can discriminate real from unreal optic flow when 2D global features and local motion signals are precisely matched. In Experiment 1, we examined whether cortical representations of motion coherence for expansion and contraction are differentially modulated by ecological validity. In Experiment 2, we further tested whether cortical regions identified in Experiment 1 could directly discriminate real from unreal optic flow, thereby probing neural sensitivity to ecological validity itself. Together, these experiments address the long-standing conflation between simplified radial motion and real optic flow, and aim to clarify how ecological structure shapes cortical representations of self-movement in the human brain.

## Results

### Experiment 1

#### Motion pattern detection task

The left two panels in **Figure 1b** plot the mean proportion of “yes” responses in the motion pattern detection task of Experiment 1 (n = 30) along with the best fitting curves against motion coherence level for the two type of motion stimuli (real and unreal optic flow). The fitted curve and the majority of data points for contraction patterns were higher than those for expansion patterns for the unreal optic flow stimuli, but this trend was reversed for the real optic flow stimuli.

**Figure 1.**
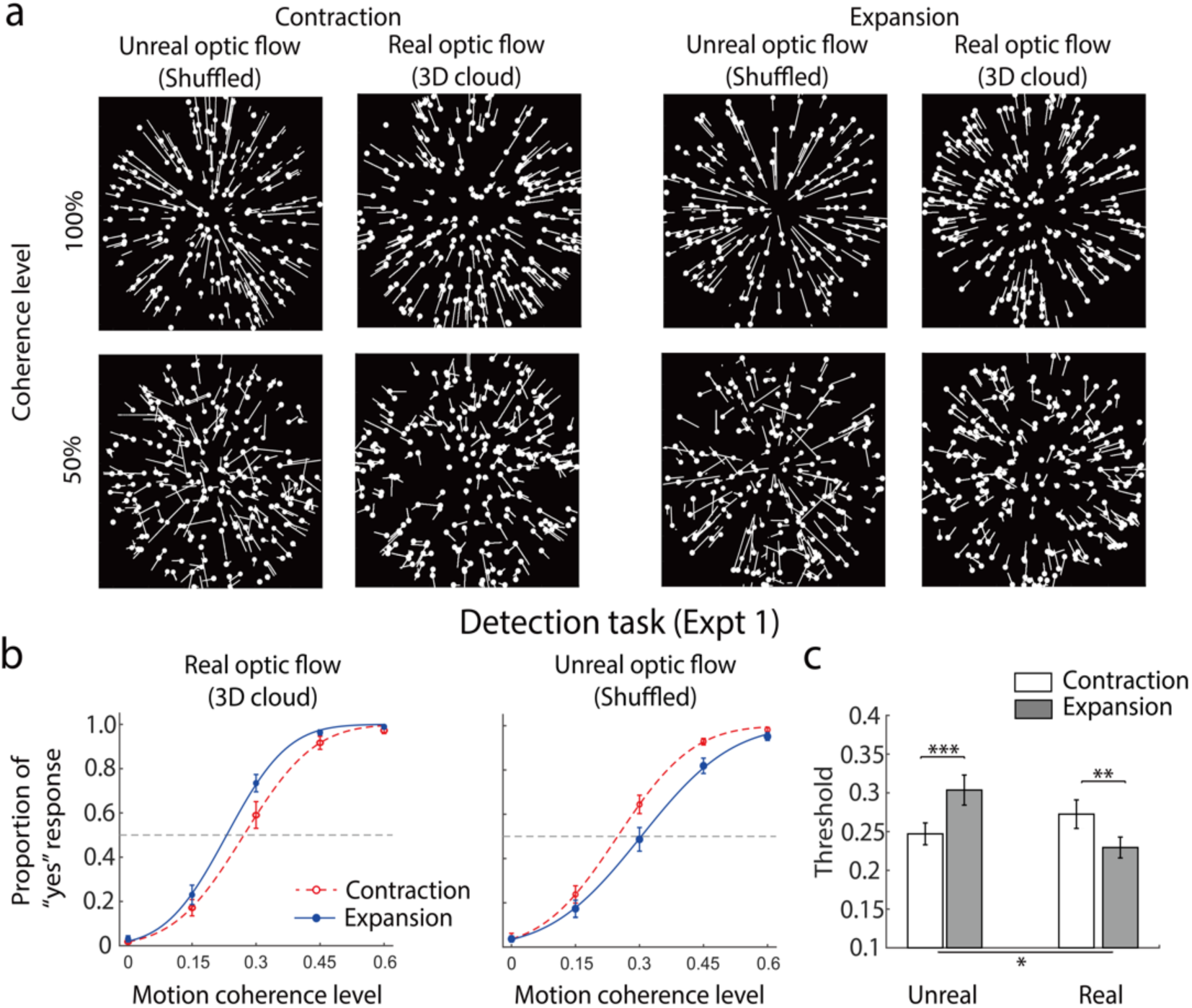
**(a)** Illustrations of the velocity fields of the unreal (shuffled radial motion) and real (3D-cloud) optic flow stimuli at 100% (upper panels) and 50% (lower panels) coherence levels in Experiment 1 and 2. The white dots indicate the dot position at the end of a trial. The white lines represent the dot motion trajectories in the trial. **(b)** Mean proportion of “yes” response data from the motion pattern detection task in Experiment 1 averaged across 30 participants along with the best fitting psychometric curves for the real (left panel) and unreal optic flow stimuli (right panel). **(c)** Mean detection threshold of the fitted curve to each participant’s data averaged across participants. Error bars indicate ±1 SE. *: p < 0.05, **: p < 0.01, ***: p < 0.001.

**Figure 1c** plots the mean 50% detection threshold of the fitted curve to each participant’s data averaged across participants. A 2 (stimulus type: real and unreal optic flow) × 2 (motion pattern: expansion and contraction) repeated-measures ANOVA on the detection threshold revealed that both the main effect of stimulus type and the interaction effect of stimulus type and motion pattern were significant (*F*(1,29) = 6.99, *p* = 0.013, *η^2^*= 0.19 and *F*(1,29) = 33.67, *p* < 0.001, *η^2^* = 0.54, respectively). The main effect of the motion pattern was not significant (*p* = 0.46). Tukey HSD tests revealed that the detection threshold was significantly lower for contraction than expansion patterns for the unreal optic flow stimuli (mean ± standard error [SE]: 0.25 ± 0.010 vs. 0.30 ± 0.014, *p* < 0.001), but this trend was reversed for the real optic flow stimuli, i.e., the detection threshold was significantly higher for contraction than expansion patterns (0.27 ± 0.013 vs. 0.23 ± 0.0095, *p* = 0.0071). This indicates that participants were more sensitive to detect contraction than expansion for unreal optic flow, but more sensitive to detect expansion than contraction for real optic flow.

#### fMRI results

To investigate the neural correlates of the behavioral differences observed in the motion pattern detection task, we conducted an fMRI experiment using a block design and the same stimulus set. The same participants with the behavioral experiment completed four scanning sessions, each containing eight runs, corresponding to the four experimental conditions defined by stimulus type (real vs. unreal optic flow) and motion pattern (contraction vs. expansion). During scanning, participants viewed the motion stimuli while performing a task-irrelevant fixation point color-change detection task. For each participant, we used standard localizer scans to define visual regions of interest (ROIs), including regions previously reported to respond to optic flow (e.g., V1, V2, V3d, V3a, V3b/KO, MT, MST, V6, V7, VIP, CSv, Pc).

#### GLM results

To identify cortical regions activated by high-coherence (60%) motion relative to a fixation baseline, we conducted group-level General Linear Model (GLM) analyses separately for each experimental condition. Following prior work (Kuai et al., 2020), ROIs were grouped into early visual (V1, V2), dorsal visual (V3d, V3a, V7, V3b/KO), dorsal motion (MT, MST), and other optic flow-related regions (VIP, V6, CSv, Pc). Whole-brain analyses revealed robust activations in early visual (V1, V2), dorsal motion (MT, MST), and dorsal visual (V3d, V3a, V7, V3b/KO) regions across all conditions (**Figure 2a**). In contrast, activations in other optic flow-related regions (V6, VIP, CSv, Pc) were relatively weak or negligible.

**Figure 2.**
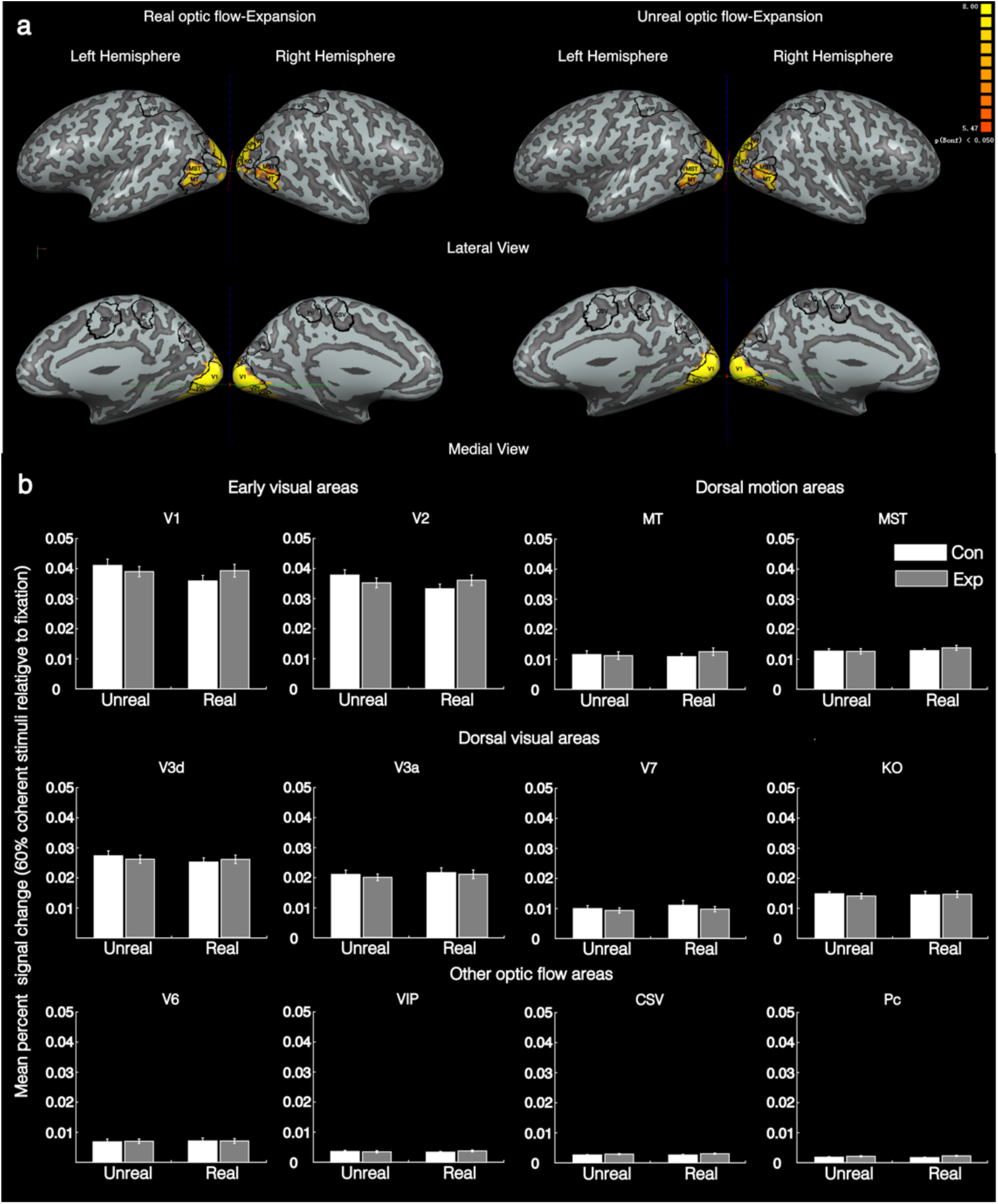
**(a)** Whole-brain multi-subject (n = 30) activation maps showing activation for the “60% coherent motion stimuli > fixation baseline” contrast on one individual inflated surface. The panels show the lateral (upper row) and medial (bottom row) views of the left and right hemispheres for the real-expansion (left panels) and unreal-expansion conditions (right panels), respectively. Clusters were considered significant if they exceeded 25 contiguous voxels at a threshold of *p* < 0.05, Bonferroni corrected, and were colored red-to-yellow. The color scale from red to yellow represents the *t*-values from the one-sample *t*-tests comparing responses of the 60% coherent motion to baseline responses. The black outlines with black texts indicate the individual functional ROIs identified from a selected participant. **(b)** Mean BOLD percent signal changes averaged across participants (across the two hemispheres) for the contrast (60% coherent stimuli - fixation baseline) against the unreal (left) and real (right) optic flow stimuli for contraction (white bars) and expansion (gray bars) patterns for each ROI. Error bars indicate ±1 SE across 30 participants.

To test whether BOLD response patterns corresponded to the behavioral effects from the motion pattern detection task, from each ROI, we selected the 500 voxels showing the strongest responses to motion. We then computed percent BOLD signal change for these voxels and averaged across the 30 participants. **Figure 2b** shows the mean percent BOLD signal change for 60%-coherence motion relative to fixation, plotted separately for real and unreal optic flow. For each ROI group, we performed a 2 (stimulus type: real vs. unreal) × 2 (motion pattern: expansion vs. contraction) × ROIs repeated-measures ANOVA. A significant interaction of stimulus type and motion pattern was observed only in the early visual group (V1, V2), *F*(1, 29) = 4.44, *p* = 0.044, *η²* = 0.13. However, Tukey HSD post hoc tests revealed no significant pairwise differences within V1 or V2 (*p*s > 0.058). In the other three ROI groups, neither the main effects of stimulus type or motion pattern (*p*s > 0.15 and *p*s > 0.33, respectively) nor their interaction (*p*s > 0.24) reached significance.

Overall, at 60% coherence, most ROIs showed similar activation for expansion and contraction under both real and unreal optic flow, suggesting that mean activation levels may be insufficient to capture differences in how motion patterns are represented across conditions. We therefore turn to MVPA to test whether the ecological validity of optic flow modulates neural representations of motion patterns.

#### MVPA results

Although univariate analyses of percent BOLD signal change did not reveal significant differences between expansion and contraction under either real or unreal optic flow in any region, such analyses may not be sensitive to distributed response patterns that differentiate experimental conditions. Given that MVPA can detect subtle but consistent differences in spatial activation patterns across voxels, we therefore applied MVPA to further examine whether ecological validity modulates neural representations of motion patterns. Specifically, we decoded coherent motion (60%, 45%, and 15% coherence) versus incoherent motion (0% coherence) within each ROI. For classification, we selected 500 voxels from each ROI, a number chosen based on prior evaluations to optimize decoding stability and ensure comparable signal-to-noise ratios across regions (see Supplementary Materials for details).

**Figure 3a** shows decoding accuracy as a function of motion coherence level, plotted separately for each ROI and experimental condition (real vs. unreal × expansion vs. contraction) in Experiment 1. To determine whether decoding accuracy exceeded chance level, we compared decoding performance against baseline (chance) for each ROI group and experimental condition. At medium to high coherence levels (30%, 45%, and 60%), decoding accuracy was significantly above baseline in all ROI groups, including early visual areas (V1/V2; *F*(1, 29) > 42.8, *p*s < 0.001, *η²* > 0.60), dorsal motion areas (MT/MST; *F*(1, 29) > 29.90, *p*s < 0.001, *η²* > 0.51), dorsal visual areas (V3d/V3a/V7/KO; *F*(1, 29) > 61.70, *p*s < 0.001, *η²* > 0.68), and other optic flow-related regions (V6/VIP/CSv/Pc; *F*(1, 29) > 20.98, *p*s < 0.001, *η²* > 0.42). At the lowest coherence level (15%), decoding performance remained above baseline only in a subset of conditions and regions. Overall, these results indicate that, across the cortical network examined, coherent motion could be reliably distinguished from random motion at moderate to high coherence levels. We next examined whether decoding accuracy increased with motion coherence. A one-way repeated-measures ANOVA with coherence level (15%, 30%, 45%, 60%) as a within-subject factor revealed a significant main effect of coherence in all ROIs and experimental conditions (*F*(3, 87) > 5.91, *p*s < 0.001, *η²* > 0.17). Tukey HSD post hoc tests showed that decoding accuracy at 45% and 60% coherence was significantly higher than at 15% coherence across all regions. These findings confirm that BOLD response patterns in all ROIs were modulated by motion coherence, for both expansion and contraction patterns and for both real and unreal optic flow.

**Figure 3.**
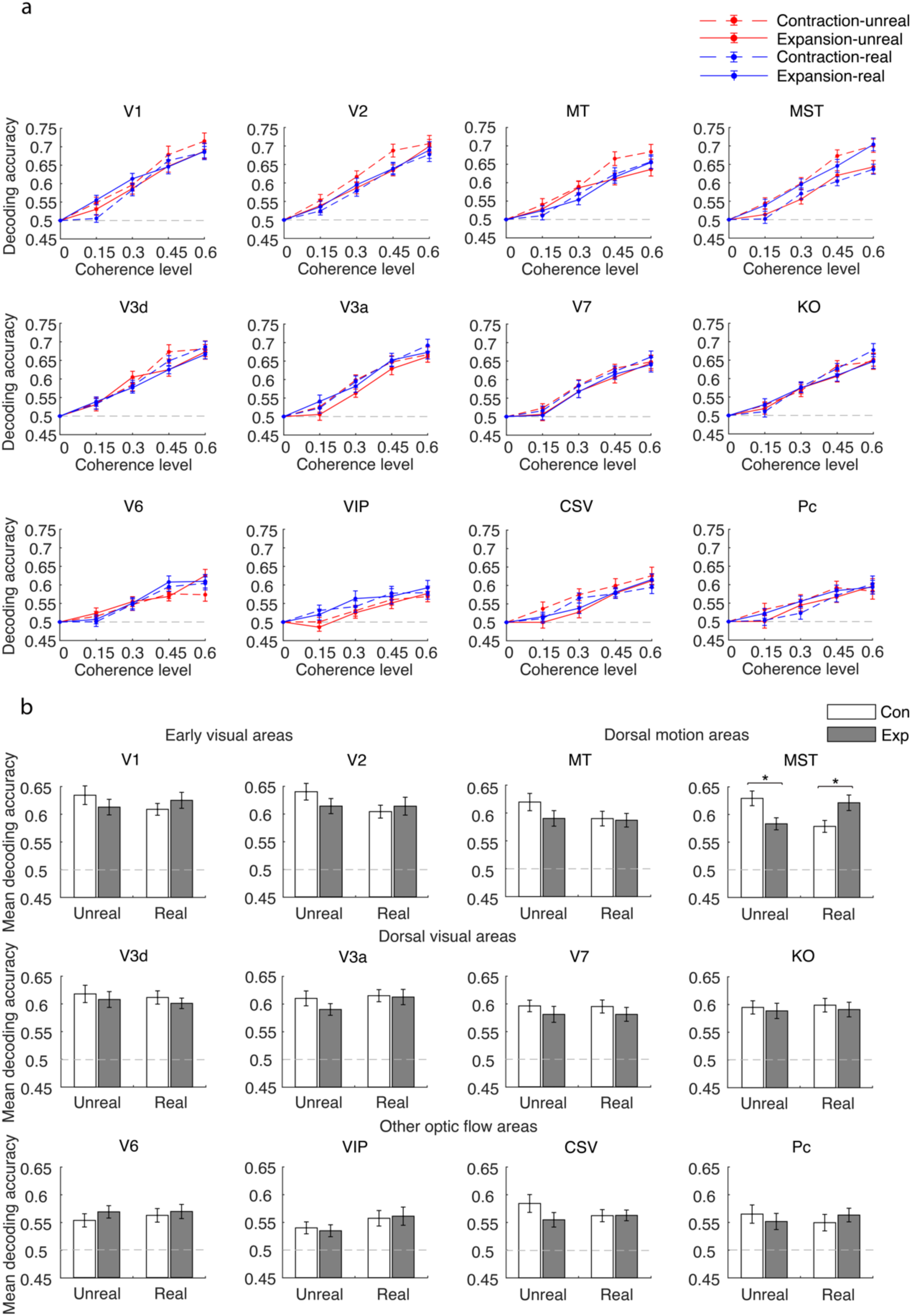
**(a)** Mean decoding accuracy for discriminating coherent versus incoherent motion of the classifier averaged across 30 participants as a function of motion coherence levels for the two types of motion stimulus in each ROI (red: unreal optic flow; blue: real optic flow; solid: expansion; dotted: contraction). The gray dashed line represents the classifier’s baseline decoding accuracy. **(b)** Mean decoding accuracy averaged across all non-zero motion coherence levels against unreal (left) and real optic flow (right) for contraction (white bars) and expansion (gray bars) patterns in each ROI. The light gray dashed lines represent the mean classifier’s baseline decoding accuracy averaged across all non-zero motion coherence levels. Error bars indicate ±1 SE across 30 participants. *: *p* < 0.05.

Having established robust decoding accuracy across regions, we then asked whether any ROI showed response patterns that mirrored the behavioral dissociation between expansion and contraction observed in the motion pattern detection task. To achieve this, we compared decoding accuracy across stimulus type (real vs. unreal optic flow) and motion pattern (expansion vs. contraction), averaging decoding performance across non-zero coherence levels (**Figure 3b**). A 2 × 2 repeated-measures ANOVA revealed a significant interaction between stimulus type and motion pattern only in MST (*F*(1, 29) = 20.52, *p* < 0.001, *η²* = 0.41). Tukey HSD post hoc tests further showed that, under unreal optic flow, decoding accuracy was higher for contraction than expansion, whereas under real optic flow this preference was reversed. No other ROI showed a comparable interaction or significant main effects (all *p*s > 0.067). These results indicate that, among the regions examined, MST uniquely showed neural response patterns that represented expansion and contraction in a manner dependent on the ecological validity of optic flow.

#### The 50% threshold of fMR-metric function

To characterize neural sensitivity in a manner comparable to behavioral detection thresholds, we derived fMR-metric functions by fitting cumulative Gaussian functions to decoding accuracy as a function of motion coherence (**Figure 4a**; see Materials and Methods). The 50% point of the fitted function was regarded as the neural threshold.

**Figure 4.**
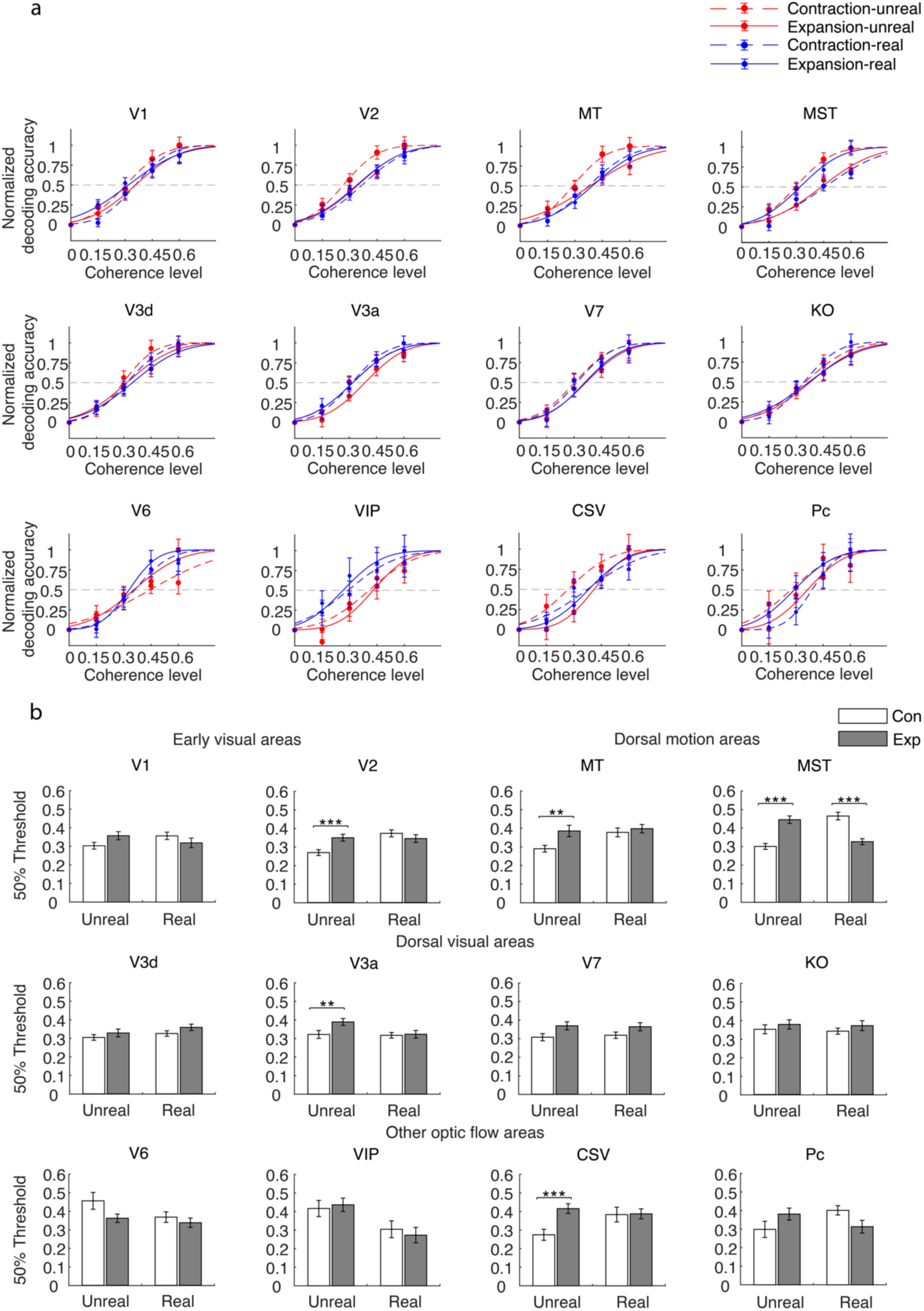
**(a)** Mean normalized decoding accuracy averaged across participants along with the best fitting cumulative Gaussian curves for the two types of stimuli (real and unreal optic flow). The gray dashed line indicates 50% normalized decoding accuracy. Error bars indicate ±1 SE across 30 participants. **(b)** Mean threshold obtained from the fMR-metric curve fitted to each participant’s data averaged across participants for the two types of motion stimuli. Error bars indicate ±1 SD of the threshold distribution, generated by bootstrapping the fitting of the cumulative Gaussian function 10,000 times. This bootstrapping method was performed by resampling (with replacement) and normalizing participants’ mean decoding accuracy data (mean ± SE) on each iteration. **: *p* < 0.01, ***: *p* < 0.001.

**Figure 4b** shows the mean 50% thresholds for expansion and contraction under real and unreal optic flow. For unreal optic flow, significantly lower thresholds were observed for contraction than expansion in V2 (mean ± SD: 0.27 ± 0.015 vs. 0.35 ± 0.018, *p* < 0.001), MT (0.29 ± 0.018 vs. 0.39 ± 0.031, *p* = 0.0019), V3a (0.32 ± 0.021 vs. 0.39 ± 0.019, *p* = 0.010), CSv (0.28 ± 0.030 vs. 0.42 ± 0.026, *p* < 0.001), and MST (0.30 ± 0.016 vs. 0.45 ± 0.020, *p* < 0.001), indicating higher neural sensitivity to contraction than expansion under unreal optic flow. By contrast, under real optic flow, only MST exhibited a significantly lower threshold for expansion than contraction (0.33 ± 0.016 vs. 0.46 ± 0.020, *p* < 0.001), whereas no other ROI showed a significant difference between motion patterns (*p*s > 0.044). These results indicate that neural sensitivity patterns in MST closely mirrored the same group of participants’ behavioral performance: higher neural sensitivity to contraction than expansion patterns for unreal optic flow, and this trend was reversed for real optic flow.

Although these results suggest that MST’s representations are modulated by ecological validity, whether MST or other optic flow-related areas can directly discriminate real from unreal optic flow.

### Experiment 2

To further validate the findings from Experiment 1 and address two key limitations, we designed Experiment 2 as a direct extension. First, although Experiment 1 revealed differences in cortical representations associated with ecological validity through decoding of motion coherence, it did not directly test whether neural activity could distinguish real from unreal optic flow. Second, because the four experimental conditions in Experiment 1 were acquired in separate scanning sessions, voxel-level variability across sessions may have influenced decoding performance. To address these issues, Experiment 2 presented real and unreal optic flow within the same scanning session and directly decoded stimulus type (real vs. unreal) under each motion pattern. This design allowed us to directly examine the neural representation of ecological validity while minimizing potential session-related confounds.

#### Stimulus type discrimination task

In Experiment 2, before scanning, participants (n = 20) completed a stimulus-type discrimination task to assess perceptual discriminability between real and unreal optic flow. Using the same stimuli as in Experiment 1, participants judged whether each stimulus depicted real or unreal optic flow at different coherence levels (15%-90% in step of 15%).

As shown in **Figure 5a**, discrimination performance increased with motion coherence for both expansion and contraction patterns. Importantly, discrimination performance did not differ between motion patterns. The mean point of subjective equivalence (PSE) and discrimination threshold (inverse slope of the fitted psychometric function) showed no significant differences between expansion and contraction (**Figure 5b**; paired *t*-tests, *<u>p</u>*s > 0.24), indicating comparable perceptual discriminability across motion patterns. Consistent with this result, d-prime values increased significantly with motion coherence for both expansion and contraction patterns (**Figure 5c**). A one-way ANOVA revealed a significant main effect of coherence level for both expansion and contraction (expansion: *F*(5, 95) = 58.34, *p* < 0.001, *η^2^* = 0.75; contraction: *F*(5, 95) = 80.52, *p* < 0.001, *η^2^* = 0.81).

**Figure 5.**
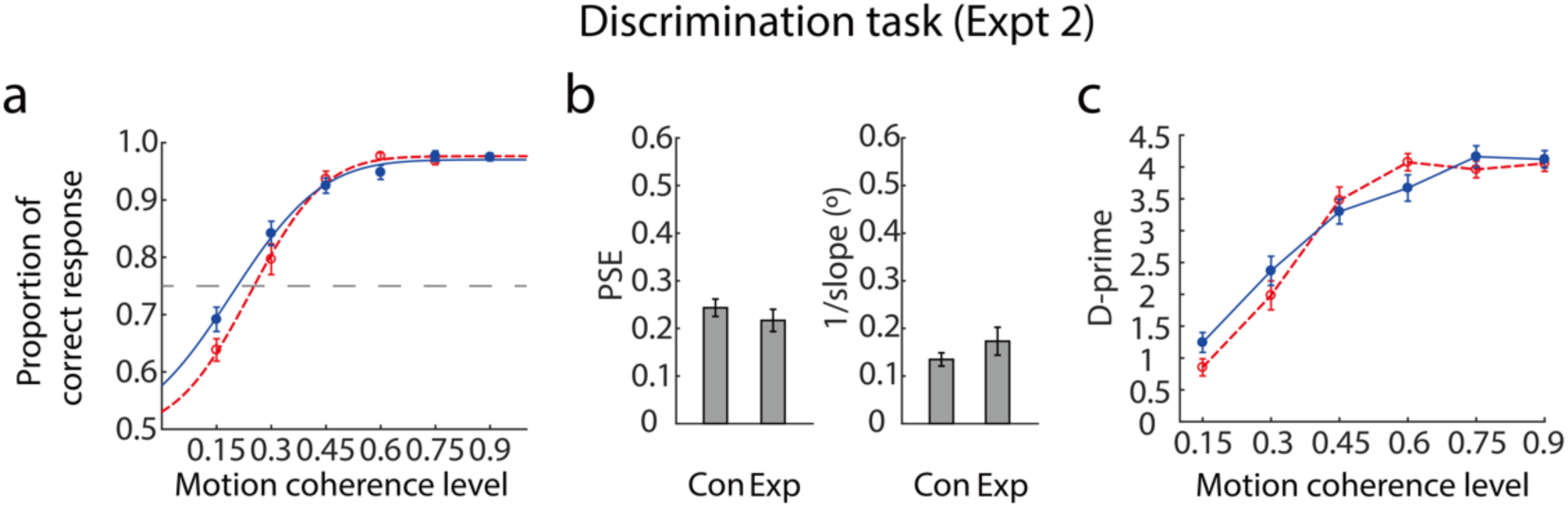
**(a)** Mean correct response data from the stimulus type discrimination task in Experiment 2 averaged across 20 participants along with the best fitting psychometric curves for the two motion patterns (expansion and contraction). (b) Mean PSE (left panel) and mean inverse of the slope (right panel) of the curves fitted to each participant’s data averaged across 20 participants against the motion patterns. (c) Mean d-prime data averaged across 20 participants as a function of motion coherence level for the two motion patterns. Error bars indicate ±1 SE.

Together, these behavioral results indicate that observers can reliably discriminate real from unreal optic flow, and that this discriminability improves with motion coherence but is not modulated by expansion or contraction. This ensures that any neural differences observed in Experiment 2 cannot be attributed to differences in behavioral task difficulty across motion patterns.

#### MVPA results

We next conducted a follow-up fMRI experiment (n = 20) using the same stimuli and a similar block design as in Experiment 1. Each participant completed two scanning sessions, one for each motion pattern (contraction or expansion), with eight runs per session. Within each session, stimulus blocks were randomly showed at three coherence levels (0%, 30%, 60%). Participants performed a task-irrelevant fixation point color-change detection task. ROIs were functionally localized in each participant and included regions previously implicated in optic flow processing (V3b/KO, MT, MST, V6, VIP, CSv, and Pc). We applied MVPA within each ROI to decode stimulus type (real vs. unreal) separately for contraction and expansion patterns.

**Figure 6** illustrates mean decoding accuracy (n = 20) for discriminating real from unreal optic flow as a function of motion coherence for each ROI and motion pattern. To determine whether decoding performance exceeded chance level, we compared decoding accuracy at each coherence level against baseline (0.5) using paired *t*-tests for each region. In MST, decoding accuracy at 60% coherence was significantly above baseline for both expansion (*t*(19) = 4.49, *p* < 0.001) and contraction patterns (*t*(19) = 2.94, *p* = 0.0083). In V6, above-baseline decoding accuracy was observed for the contraction pattern at 60% coherence (*t*(19) = 2.56, *p* = 0.019). At 30% coherence, above-baseline decoding accuracy was found in V3b/KO for the contraction pattern (*t*(19) = 2.39, *p* = 0.027) and in V6 for the expansion pattern (*t*(19) = 2.29, *p* = 0.019). No other ROI showed decoding accuracy significantly above baseline (*p*s > 0.063).

**Figure 6.**
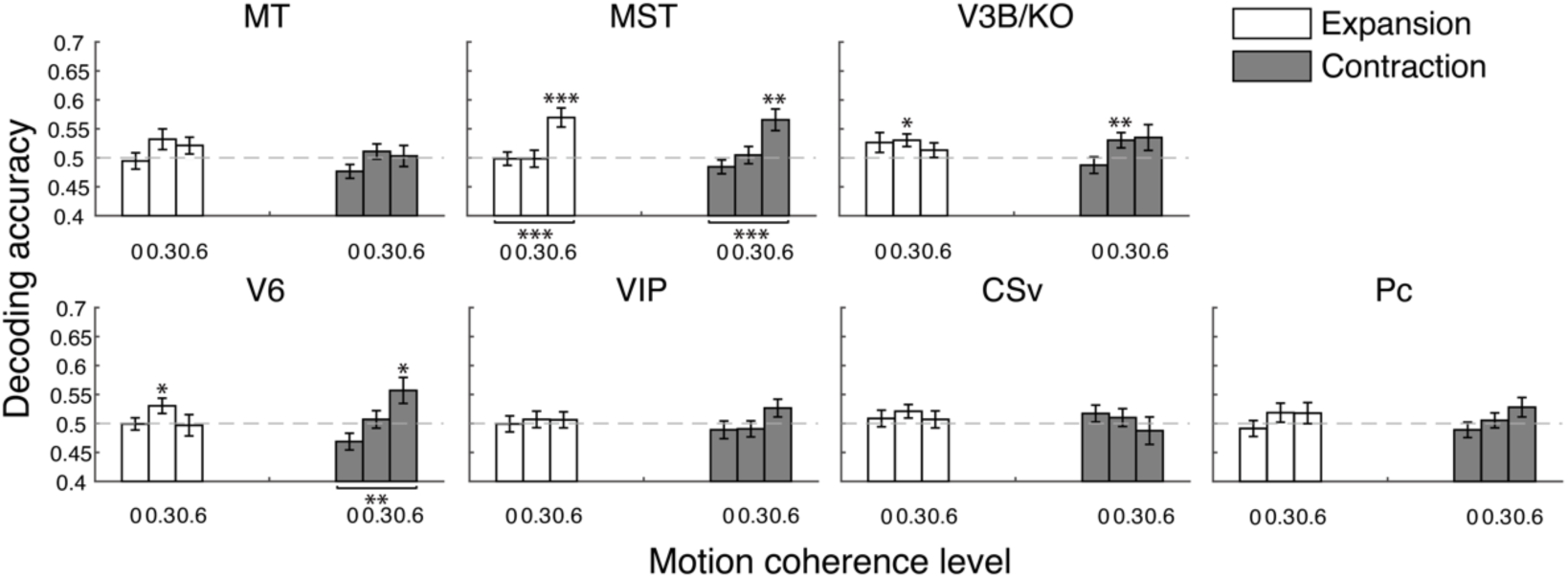
**(a)** Mean decoding accuracy for discriminating real from unreal optic flow stimuli of the classifier averaged across 20 participants as a function of motion coherence levels for expansion (white bars) and contraction patterns (gray bars) in each ROI. The gray dashed line represents the mean classifier’s baseline decoding accuracy averaged across all motion coherence levels. Error bars indicate ±1 SE across 20 participants. *: *p* < 0.05, **: *p* < 0.01, ***: *p* < 0.001.

To assess whether decoding accuracy varied systematically with coherence level, we conducted a one-way repeated-measures ANOVA with coherence level as a within-subject factor for each ROI and motion pattern. This analysis revealed a significant main effect of coherence level in MST for both expansion (*F*(2, 38) = 10.99, *p* < 0.001, *η²* = 0.37) and contraction patterns (*F*(2, 38) = 7.64, *p* = 0.0016, *η²* = 0.29). Post hoc Tukey HSD tests showed that decoding accuracy at 60% coherence was significantly higher than at both 30% and 0% coherence for both motion patterns (expansion: 0.58 ± 0.017 vs. 0.50 ± 0.015 and 0.49 ± 0.013; contraction: 0.56 ± 0.020 vs. 0.50 ± 0.015 and 0.48 ± 0.013; *p*s < 0.033). A significant main effect was also observed in V6 for the contraction pattern (*F*(2, 38) = 6.56, *p* = 0.0036, *η²* = 0.26), with decoding accuracy at 60% coherence significantly exceeding that at 0% coherence (0.56 ± 0.022 vs. 0.47 ± 0.015; *p* = 0.0026).

## Discussion

The present study combined behavioral measures with multivariate fMRI analyses across two experiments to examine how the human visual system processes optic flow that varies in ecological validity. Across experiments, we found convergent evidence that the dorsal visual area MST not only distinguishes real optic flow from unreal variants, but also systematically alters its sensitivity to expansion and contraction patterns depending on whether the stimulus preserves the natural 3D structure of self-movement. These results demonstrate that MST does not operate with a fixed preference for a particular motion pattern, but flexibly adjusts its response profile according to the ecological structure of the visual input.

A key finding of the present study is that MST responses depend critically on the presence of a continuous speed gradient consistent with natural self-movement. During forward locomotion, optic flow forms a radial expansion pattern in which image speed increases with eccentricity, reflecting the geometry of a rigid 3D environment and providing essential information about direction and velocity of self-movement (Cutting et al., 1992; Gibson, 1966; Rogers & Graham, 1979). In the present experiments, only real optic flow preserved this ecologically valid relationship between speed and depth. Under these conditions, both behavioral sensitivity and neural sensitivity in MST were higher for expansion than contraction patterns. When the same motion patterns were presented as unreal optic flow that disrupted this natural structure, this preference reversed. These findings indicate that MST responses are determined not by motion pattern alone, but by whether the global speed structure of the stimulus corresponds to ecologically valid self-movement.

Neurophysiological evidence from non-human primates supports this interpretation. The human MST area is widely regarded as the putative homologue of the dorsal subdivision of macaque MST (MSTd), which plays a central role in processing optic flow and self-movement cues along the dorsal visual pathway (Dukelow et al., 2001; Huk et al., 2002). Studies using unreal optic flow stimuli have reported relatively little sensitivity in MSTd neurons to speed gradients that do not reflect natural 3D structure (Orban et al., 1995; Tanaka et al., 1989). In contrast, studies using real optic flow that simulated self-movement through a rigid environment demonstrated strong neuronal preferences for gradients consistent with natural locomotion and sensitivity to motion parallax (Duffy & Wurtz, 1997; Upadhyay et al., 2000). Together, these findings suggest that MSTd neurons selectively encode speed gradients that convey meaningful information about 3D scene structure and self-movement rather than unreal variations in speed distribution.

The present fMRI results extend these principles to the human visual system. In Experiment 1, MST exhibited a systematic reversal of motion pattern sensitivity depending on ecological validity, with an expansion advantage under real optic flow and a contraction advantage under unreal optic flow. Behavioral performance showed a highly similar pattern, indicating a close correspondence between perceptual sensitivity and neural representations in MST. Experiment 2 further demonstrated that MST reliably discriminates real from unreal optic flow independent of motion pattern when the global continuity of the speed gradient is preserved. When motion coherence was reduced to 0% and the global structure of the flow field was disrupted, MST no longer distinguished between the two stimulus types, despite the presence of local speed differences. This finding demonstrates that MST responses depend on coherent global motion structure rather than isolated local motion cues.

### Contraction bias as a default mode of motion processing under unreal optic flow

When optic flow lacks coherent self-movement structure, the visual system shows a reliable bias toward contraction, as observed in the unreal optic flow conditions of the present study. Behaviorally, contraction patterns were detected more easily than expansion, and this advantage was mirrored by higher neural sensitivity in MST and other vision- or motion-related regions, including V2, MT, V3a, and CSv. Similar contraction biases have been consistently reported in psychophysical, neuroimaging, and electrophysiological studies using unreal optic flow stimuli in which dots move at constant speed or lack natural depth structure (Edwards & Badcock, 1993; Edwards & Ibbotson, 2007; Giaschi et al., 2007; Shirai et al., 2009; Snowden & Milne, 1996).

One explanation is that, in the absence of reliable self-movement cues, the visual system relies on more general motion processing mechanisms that are optimized for central viewing. Contraction produces inward-converging motion vectors toward the fovea, resulting in higher local directional coherence and more efficient signal integration within a limited spatial window. This advantage is amplified by the non-uniform sampling properties of the visual system, with higher spatial resolution and signal-to-noise ratio in central vision compared with the periphery (Curcio et al., 1990; Curcio & Allen, 1990; Watson, 2014). Under fixation, centripetal motion therefore benefits disproportionately from foveal-centered integration, leading to enhanced detectability and stronger neural responses. Consistent with this account, psychophysical studies have shown that contraction is more easily detected than expansion at low eccentricities, with the advantage diminishing at larger eccentricities (Edwards & Badcock, 1993). Together, these observations suggest that contraction bias under unreal optic flow reflects a default mode of motion processing that emphasizes local coherence and foveal-centered integration.

### Experience-dependent emergence of expansion bias under real optic flow

Against this default mode, the expansion advantage observed under real optic flow can be interpreted as reflecting a reweighting of motion processing driven by ecological structure and long-term experience. In natural vision, forward self-movement occurs far more frequently than backward movement, making expanding optic flow the statistically dominant pattern during natural forward locomotion. Real optic flow preserves the key self-movement cues that allow expansion to serve as a reliable signal for heading and self-movement estimation (Cutting et al., 1992; Gibson, 1966; Rogers & Graham, 1979).

According to efficient coding theory, neural systems allocate greater representational resources to sensory patterns that are frequently encountered and behaviorally relevant (Attneave, 1954; Barlow, 1961). Consistent with this framework, repeated exposure to specific motion statistics enhances perceptual sensitivity and induces experience-dependent changes in neuronal tuning throughout the visual system (Dragoi et al., 2000; Iliescu, 2022; Sur et al., 2002; Watanabe et al., 2001; Zhang et al., 2010). Within this theoretical framework, the predominance of forward self-movement during everyday locomotion may gradually bias areas in the dorsal stream such as MST toward more efficient processing of expanding motion patterns when the stimulus supports a self-movement interpretation.

Evidence for such experience-dependent tuning also comes from electrophysiological and developmental studies. Several EEG studies have identified the N2 component as an index of motion processing efficiency, with generators localized to motion-sensitive cortical regions such as MT+ (Ahlfors et al., 1999; Probst et al., 1993). Using real optic flow stimuli, Vilhelmsen et al. (2019) reported longer N2 latencies for backward than forward self-movement at high speeds, indicating less efficient processing of the less familiar backward motion. Importantly, this asymmetry was present only in infants with self-initiated locomotor experience and was absent in younger infants without such experience, suggesting that sensitivity to forward expansion emerges through active interaction with the environment (Mukherjee et al., 2002). Developmental evidence further shows that neural responses to forward optic flow become progressively faster with age, consistent with increasing specialization for expansion patterns as visual and motor systems mature (Rasulo et al., 2021).

Together, these findings suggest that the expansion advantage observed under real optic flow does not reflect an intrinsic preference for expansion per se, but rather the outcome of long-term adaptation to the statistical regularities of natural self-movement. When optic flow preserves coherent 3D self-movement structure, the visual system shifts away from default foveal-centered motion integration and engages experience-tuned mechanisms optimized for self-movement perception.

### Ecological validity as a continuum

Although the present study operationally categorized optic flow stimuli as “real” versus “unreal”, converging evidence from prior work suggests that the ecological validity of optic flow is more accurately described as a continuum, rather than a strict categorical distinction. Across neurophysiological, neuroimaging, and psychophysical studies, the critical determinant of motion pattern sensitivity appears to be the extent to which a stimulus preserves coherent 3D self-movement cues, such as continuous speed gradients, motion parallax, and scene rigidity. Highly naturalistic optic flow is typically associated with an expansion advantage in MSTd (Duffy & Wurtz, 1995; Gu et al., 2010) and more efficient neural processing of forward self-movement in humans (Vilhelmsen et al., 2019). Importantly, some studies using unreal optic flow with linearly increasing radial speed have also reported an expansion bias, likely because such stimuli retain a coherent global speed gradient and evoke a percept of motion in depth (Duffy & Wurtz, 1991a; Geesaman & Andersen, 1996; Graziano et al., 1994; Heuer & Britten, 2004; Tanaka & Saito, 1989). By contrast, when the ecological structure of optic flow is further degraded, for example by removing speed gradients altogether, both behavioral and neural measures often reveal a contraction advantage (Edwards & Badcock, 1993; Edwards & Ibbotson, 2007; Giaschi et al., 2007; Shirai et al., 2009; Snowden & Milne, 1996).

Seen in this light, the present findings are best understood within a graded framework in which motion pattern sensitivity in MST reflects the degree to which optic flow preserves coherent global self-movement structure. The expansion advantage observed under real optic flow in the present study corresponds to a higher level of ecological validity, whereas the contraction bias under unreal optic flow reflects a shift toward lower ecological validity when global self-movement cues are disrupted.

### Network-level sensitivity to global motion coherence

Following these findings, it is important to consider the broader network context in which MST operates. Across all visual and optic flow-responsive regions examined, including early visual cortex and higher-level dorsal stream areas, neural responses were robustly modulated by motion coherence under all stimulus conditions.

Decoding performance reliably distinguished coherent from random motion across regions, consistent with extensive prior neuroimaging univariate and multivariate evidence that global radial motion coherence engages a distributed visual network (Braddick et al., 2001; Cowey et al., 2006; Fattori et al., 2009; Koyama et al., 2005; Pitzalis et al., 2010; Wada et al., 2016).

These results indicate that the contraction bias does not reflect a reduced sensitivity to global motion per se. Instead, it suggests that ecological structure selectively modulates how global motion information is weighted and represented within this network, with MST showing the clearest and most behaviorally relevant sensitivity to changes in self-movement structure.

### Relation to previous findings using highly naturalistic stimuli

In contrast to the present findings, Di Marco et al. (2021) reported stronger BOLD responses to forward than backward self-movement in regions such as V3A, V6, CSv, and Pc when using highly naturalistic virtual scenes. The absence of comparable expansion advantages in these regions in the current study likely reflects differences in stimulus design rather than fundamental discrepancies in neural mechanisms. Di Marco et al. employed immersive environments containing explicit ground planes, sky, and scene layout, which may more strongly engage large-scale spatial representations and multisensory self-motion processing. By comparison, the real optic flow used in the present study was intentionally designed as a controlled 3D dot cloud that preserved essential global motion structure while minimizing higher-level scene semantics. This approach allowed us to isolate the contribution of global motion structure to optic flow processing and revealed MST as the region most consistently sensitive to ecological validity. Future work that parametrically varies scene realism while preserving global motion structure will be essential for clarifying how different levels of ecological validity shape the distribution of self-movement representations across the dorsal visual system.

The present study focused on radial optic flow because it provides direct information about heading and self-movement, making it particularly well suited for examining how ecological validity shapes motion processing. Converging evidence from non-human primate neurophysiology and human neuroimaging shows that MST is especially responsive to radial motion components compared with other motion types (Duffy & Wurtz, 1991a, 1991b; Graziano et al., 1994; Pitzalis et al., 2013). By using expanding and contracting radial motion, the present design enabled a targeted examination of how ecologically valid self-movement structure reshapes motion pattern sensitivity within the dorsal visual stream. Whether similar ecological reweighting effects extend to other motion types, such as spiral or rotational flow, remains an open question and will require future studies that systematically vary motion structure while controlling ecological validity.

Together, the present findings demonstrate that motion pattern sensitivity in human MST is not a fixed property tied to expansion or contraction per se, but a flexible outcome shaped by the ecological structure of optic flow and long-term experience with self-movement. When optic flow preserves coherent 3D structure and supports self-movement perception, MST exhibits an expansion advantage consistent with natural forward locomotion. When such structure is absent, motion processing reverts to a default mode that favors contraction through local, foveal-centered integration. This pattern reconciles previously mixed findings across studies using different optic flow stimuli and highlights ecological validity as a critical factor in interpreting neural responses to motion. More broadly, these results suggest that MST serves as a key site where global motion signals are dynamically reweighted according to their relevance for self-movement, providing a neural bridge between basic motion processing and ecologically grounded perception.

## Materials and Methods

### Participants

Thirty college students and staff (29 naive to the specific purpose of the study) between the ages of 19 and 27 years (14 males, 16 females; mean age ± SD: 23.4 ± 2.1 yrs) participated in Experiment 1, and seven of them also participated in Experiment 2. Twenty participants (19 naive to the specific purpose of the study) between the ages of 20 and 28 years (11 males, 9 females; mean age ± SD: 23.5 ± 2.2 yrs) participated in Experiment 2. All participants had normal or corrected-to-normal vision. We obtained written informed consent from all participants before the commencement of the experiment. The consent form was approved by the Institutional Review Board at New York University Shanghai. The determination of the sample size was based on relevant previous studies.

### Stimuli

The visual stimuli were generated using the same procedures as in Experiment 2 of the previous study (Li et al., 2025). Two types of optic flow stimuli were used in both experiments: real optic flow and unreal optic flow. Real optic flow was generated by simulating observer translation through a rigid 3D-dot cloud, producing expanding or contracting radial flow with motion-parallax cues. Unreal optic flow was generated from the corresponding real optic flow displays by shuffling dots’ image velocities while preserving their initial positions and motion directions, thereby matching the two stimulus types in 2D global structure and local image motion signals but disrupting coherent self-movement structure.

All displays consisted of 145 white dots (0.09° diameter; 98.7% luminance contrast) presented within a 19° aperture, with dot motion removed within a central 1° region to minimize position cues. For the real-optic-flow stimuli, dots were distributed within a 3D volume (depth range: 1.5–5.3 m). Forward and backward scene speeds (2.09 m/s and 1.72 m/s) were chosen to equate the modal image speed between expansion and contraction (5.50°/s). When dots exited the aperture, the same number of dots was regenerated according to the depth distribution for each stimulus type to keep dot number constant. Dot placement followed Alliston (2004) to ensure a uniform dot distribution on the image plane. Additional implementation details, including stimulus generation and validation procedures, are provided in Li et al. (2025).

### Procedure

Before scanning, participants completed two psychophysical tasks: a motion pattern detection task (corresponding to Experiment 1) and a stimulus type discrimination task (corresponding to Experiment 2). Both tasks were performed outside the scanner, and the motion stimulus duration was 600 ms in all experiments.

For the motion pattern detection task, four experimental conditions were tested: 2 stimulus types (real vs. unreal) × 2 motion patterns (contraction vs. expansion). In all conditions, the CoM location was at the center of the display (i.e., 0°). On each trial, a stimulus with a motion coherence level randomly chosen from 0% to 60% (in steps of 15%) was presented. The motion coherence level indicates the percentage of signal dots in the stimulus. The rest were noise dots generated by randomly perturbing the image motion direction (in the range of 0° to 359°) of the dots while keeping their image motion speed and acceleration intact. Accordingly, the noise and the signal dots shared similar speed and acceleration profiles throughout the trial. At the end of the trial, participants were asked to indicate whether they perceived any coherent motion pattern by clicking the left (yes) or right (no) mouse button. Each of the four conditions included 30 trials per coherence level, resulting in a total of 600 trials (4 conditions × 5 coherence levels × 30 trials).

For the stimulus type discrimination task, participants viewed two experimental conditions (2 motion patterns: contraction vs. expansion), and completed a two-interval forced-choice (2IFC) task to judge whether the first or second interval contained a real optic flow stimulus. Each trial consisted of two consecutive intervals: one with a real optic flow stimulus and one with an unreal optic flow stimulus, presented in random order and separated by a 400-ms blank screen. The motion coherence level (from 15% to 90% in steps of 15%) was identical in both intervals and randomly selected on each trial. At the end of the trial, participants reported which interval contained the real optic flow by clicking the left (first interval) or right (second interval) mouse button. Each of the two motion patterns (expansion and contraction) was tested across six coherence levels, with 30 trials per level, resulting in 360 trials in total (2 motion patterns × 6 coherence levels × 30 trials).

In each task, trials were blocked by experimental conditions and randomized within each block. The testing order of experimental conditions was counterbalanced between participants. Participants were given 12 practice trials randomly selected from the experimental trials to familiarize themselves with the task before each task started. No feedback was given in the practice and experimental trials to prevent participants from developing artificial strategies that could distort their natural perceptual processes to perform the task. The motion pattern detection task lasted about 30 min and the stimulus type discrimination task lasted about 25 min.

The fMRI experiments adopted a block design and used the same stimuli as the behavioral tasks. Experiment 1 consisted of four scanning sessions, each corresponding to one of the four experimental conditions: stimulus type (real vs. unreal) × motion pattern (expansion vs. contraction). Each session included eight runs. Within each run, participants viewed 15 stimulus blocks (five motion coherence levels: 0%, 15%, 30%, 45%, 60%; each repeated three times) interleaved with 4 fixation blocks. Experiment 2 included two scanning sessions, each corresponding to one of the two motion patterns (expansion vs. contraction). Each session also had eight runs. Within each run, participants viewed 24 stimulus blocks (2 stimulus types: real vs. unreal × 3 coherence levels: 0%, 30%, 60%; each repeated four times), interleaved with 3 fixation blocks.

In both experiments, each stimulus block contained 16 1-s trials of a 600-ms motion stimulus at one motion coherence level with a red fixation point (diameter: 0.2°) in the center of the display followed by a 400-ms gray fixation point in a blank screen. The testing order of stimulus was randomized in each run. Each fixation block also had 16 1-s trials of a 600-ms display with no stimulus but a red fixation point followed by a 400-ms gray fixation point in the center of a blank screen at the beginning, middle, and end of the run. The purpose of the fixation block was to acquire baseline brain activations in each run. In 25% of trials, the color of a fixation point was changed to blue. Participants were asked to fixate on the center of the screen and perform the fixation point color change detection task by pressing a button to report the trials containing the blue fixation point. Each scanning session lasted about 1 hr 20 min and was conducted on a separate day.

In a separate session that lasted about 1 hr, we identified each participant’s ROIs related to vision and ROIs previously reported to respond to optic flow using standard functional localizer protocols. These ROIs include the early visual (V1, V2), dorsal motion (MT, MST), dorsal visual (V3d, V3a, V7 and V3b/KO) and optic flow-related areas (V6, VIP, CSv and Pc). Specifically, we identified the retinotopic visual areas (V1, V2, V3d, V3a and V7) using standard retinotopic mapping procedures with rotating wedge stimuli (DeYoe et al., 1996; Engel et al., 1994; Sereno et al., 1995), and localized V3b/KO (Dupont, 1997; Zeki et al., 2003), MT and MST (Huk et al., 2002), V6, CSv, VIP and Pc (Cardin & Smith, 2010; Wall & Smith, 2008) using independent localizers as described in the cited studies (see Supplementary Materials for details).

To confirm that different patterns of brain activations were not due to different patterns of the eye movements for different stimuli, we recorded eye movements of all participants throughout each trial during scanning using an Eyelink 1000 plus eye tracker (1k Hz, SR Research Ltd., Ontario, Canada). Analyses of fixation stability and eye-movement metrics revealed no systematic differences across conditions.

### Apparatus and data acquisition

For the motion pattern detection and stimulus type discrimination tasks, stimuli were programmed in MATLAB using Psychophysics Toolbox 3 and OpenGL. Visual stimuli were presented on an ASUS VG278H 27-inch LCD monitor (resolution: 1920 × 1080 pixels; refresh rate: 120 Hz). Participants viewed the display binocularly with their cyclopean eye aligned to the center of the screen. Head position was stabilized using a chin rest at a viewing distance of 90 cm. The testing room was light excluded.

In both fMRI experiments, stimuli were rendered using Psychtoolbox-3 and presented on a Sinorad MR-compatible LCD monitor (resolution: 1920 × 1080 pixels; refresh rate: 60 Hz) in the scanning room. Participants lay supine in a Siemens Magnetom Prisma Fit 3T MRI scanner (19 participants in Experiment 1) or a Siemens Magnetom Prisma 3T MRI scanner (11 participants in Experiment 1 and 20 participants in Experiment 2). Visual stimuli (19° diameter) were viewed binocularly through mirrors at a viewing distance of 150 cm. Head motion was minimized using a 64-channel head coil.

Functional images were acquired using echo-planar imaging (EPI) with the following parameters: voxel size = 1.5 × 1.5 × 1.5 mm, echo time (TE) = 30 ms, flip angle (FA) = 80°, matrix size = 96 × 96, field of view (FOV) = 192 × 192 mm², 60 slices acquired in ascending interleaved order (slice thickness = 2.0 mm, inter-slice gap = 0.2 mm), and repetition time (TR) = 2000 ms. A high-resolution T1-weighted anatomical image was also acquired (voxel size = 0.9 × 0.9 × 0.9 mm, TE = 2.32 ms, FA = 8°, FOV = 240 × 240 mm², 192 slices, no gap, TR = 2530 ms).

### Data analysis

#### Behavioral tasks

For the motion pattern detection task in Experiment 1, the proportion of “yes” responses was fitted as a function of motion coherence level using a cumulative Gaussian function for each participant. The motion coherence corresponding to 50% “yes” responses on the fitted curve was taken as the detection threshold, which is inversely related to perceptual sensitivity.

For the stimulus type discrimination task in Experiment 2, the proportion of correct responses was fitted as a function of motion coherence level using a cumulative Gaussian function. The point of subjective equality (PSE) was defined as the coherence level corresponding to 75% correct responses. Discrimination threshold was quantified as the inverse slope of the fitted curve. In addition, discrimination performance was assessed using d’prime values from signal detection theory, computed from hit rates (correct identification of real optic flow) and false alarm rates (incorrect identification of unreal optic flow as real).

#### fMRI preprocessing

Anatomical images were transformed into standard Montreal Neurological Institute (MNI) space and inflated using BrainVoyager QX (Brain Innovations, Maastricht, Netherlands). Functional data preprocessing included slice timing correction, 3D motion correction, linear trend removal, and temporal high-pass filtering. Functional images were coregistered to anatomical images and transformed into MNI space. All functional data were resampled into 1-mm isotropic volume time courses using nearest-neighbor interpolation without spatial smoothing.

#### GLM

We first normalized the BOLD time-course data within each run by computing Z-scores, in order to reduce run-wise baseline variability prior to subsequent averaging across stimulus blocks. Z-scores were calculated by subtracting the mean of the time course from the raw signal and dividing by its standard deviation. To account for the hemodynamic response delay in block-wise signal estimation, the time courses were shifted forward by 4 s before averaging across trials within each stimulus block.

To obtain an overview of cortical regions sensitive to coherent motion at the whole-brain level in Experiment 1, we conducted a group-level multi-subject GLM using BrainVoyager QX. Functional data from all participants were transformed into standard MNI space and concatenated at each voxel. For each experimental condition, stimulus blocks were modeled as boxcar functions convolved with a canonical hemodynamic response function (HRF) and entered as predictors in a standard GLM. One-sample *t*-tests were then performed to compare responses to 60% coherent motion against the fixation baseline. A cluster-size threshold of 25 voxels was applied, and Bonferroni correction was used to control for multiple comparisons. Statistical parametric maps were generated for the contrast of coherent motion versus fixation for each condition.

Given inter-individual variability in ROI locations, we next examined BOLD responses at the ROI level using participant-specific GLMs. For each participant and each ROI, voxel-wise responses to 60% coherent motion were estimated relative to the fixation baseline. Within each ROI, the 500 voxels showing the strongest activation were selected, following the same voxel-selection criterion used in subsequent multivariate analyses. Percent signal change was then averaged across these voxels and across participants.

#### MVPA

We performed ROI-based multivoxel pattern analysis (MVPA; Haynes & Rees, 2005; Kamitani & Tong, 2005) using MATLAB and the CoSMoMVPA toolbox (Oosterhof et al., 2016). To improve classification stability while reducing feature redundancy, voxels within each ROI were first ranked based on their overall responsiveness across experimental conditions, without reference to class labels. We then applied support vector machine recursive feature elimination with correlation bias reduction ( SVM-RFE-CBR; Yan & Zhang, 2015), which iteratively removes voxels with minimal impact on classification performance and produces a ranked list of informative features. The top-ranked voxels were selected for subsequent classification analyses.

For Experiment 1, within each ROI and experimental condition (stimulus type × motion pattern), we trained a linear support vector machine (SVM) classifier to discriminate BOLD response patterns evoked by 0% and 60% coherent motion using data from seven of the eight runs. The trained classifier was then tested on the left-out run to predict incoherent versus coherent motion at each coherence level (0%, 15%, 30%, 45%, and 60%). This procedure was repeated using leave-one--out cross-validation, and decoding accuracy at each coherence level was averaged across folds. This approach allowed us to assess whether representations learned at high coherence generalized to intermediate coherence levels.

The same MVPA procedure was applied in Experiment 2 to evaluate whether neural activity could discriminate real from unreal optic flow. For each ROI and motion pattern, a linear SVM was trained to classify real versus unreal optic flow at 60% coherence using data from seven runs and tested on the remaining run to predict stimulus type at each coherence level (0%, 30%, and 60%). Mean decoding accuracy was obtained by averaging across cross-validation folds.

To evaluate whether decoding accuracy significantly exceeded chance, we performed a permutation-based analysis in which stimulus labels in the training data were randomly shuffled. Specifically, for Experiment 1, we randomly reassigned labels for the 0% and 60% coherence stimuli; for Experiment 2, we shuffled the labels of the 60% coherent real and unreal stimuli. For each iteration, we trained the classifier on the label-shuffled training data and tested it on the corresponding stimulus blocks in the unselected run, either the 0% and each non-zero coherence level (Experiment 1) or real and unreal stimuli at each coherence level (Experiment 2). This procedure was repeated 1,000 times. The decoding accuracies from these permutations were averaged to generate a baseline (chance-level) decoding accuracy for each condition.

#### The 50% threshold of fMR-metric function

For Experiment 1, we further quantified neural sensitivity by constructing fMR-metric functions based on decoding accuracy across motion coherence levels. Specifically, for each ROI and experimental condition, we first trained a two-way linear SVM classifier to discriminate BOLD responses evoked by 0% and 60% coherent motion using stimulus blocks from randomly selected training runs. To estimate a baseline decoding performance, the trained classifier was tested on identical stimuli (0% vs. 0% coherence) by duplicating the 0% coherent stimulus blocks and applying the same prediction procedure to the testing runs. This baseline accuracy reflects classifier performance in the absence of discriminative motion information and was used to normalize decoding accuracy across ROIs and conditions.

Decoding accuracy at each coherence level was then linearly scaled by subtracting the baseline accuracy and dividing by the difference between the maximum decoding accuracy observed across all experimental conditions (real vs. unreal optic flow × expansion vs. contraction × coherence levels) and the baseline. This normalization ensured that fMR-metric functions were defined on a common scale across conditions. The normalized decoding accuracy was subsequently fitted with a cumulative Gaussian function as a function of motion coherence level for each ROI and condition, yielding fMR-metric functions (S. Li et al., 2009). The motion coherence corresponding to 50% of the normalized decoding accuracy on the fitted curve was defined as the fMR-metric threshold, providing an index of neural sensitivity analogous to behavioral detection thresholds.

To statistically compare fMR-metric thresholds between expansion and contraction patterns under real and unreal optic flow, we employed a bootstrap resampling procedure (Fischer & Whitney, 2014). Specifically, 10,000 bootstrap samples were generated by resampling with replacement from the group-level mean decoding accuracy data (mean ± SE) for each condition. For each bootstrap sample, decoding accuracy was normalized and fitted with a cumulative Gaussian function to estimate the 50% threshold as described above. This procedure yielded a distribution of threshold estimates for each of the four experimental conditions (real/unreal optic flow × expansion/contraction) within each ROI. Threshold differences between contraction and expansion were then computed separately for real and unreal optic flow. Statistical significance was assessed by determining the percentile of zero within the distribution of threshold differences. Because four within-ROI comparisons were performed, a Bonferroni correction was applied, resulting in a corrected significance threshold of *p* = 0.05/4 (0.0125).

#### Eye Movement

Eye movements were recorded during all scanning sessions to ensure that differences in neural responses were not driven by systematic differences in fixation behavior across stimulus conditions (see Supplemental Materials for details). Eye position was recorded continuously throughout each trial using an Eyelink 1000 Plus (1 kHz; SR Research Ltd., Ontario, Canada).

To characterize overall fixation stability, we computed histograms of horizontal and vertical eye positions relative to the center of the display across all participants and experimental sessions. Eye movement variability was quantified by calculating the root-mean-square error (RMSE) of horizontal and vertical eye positions relative to fixation, as well as the number and amplitude of saccades for each participant. Eye movement data were collected across all scanning sessions in both experiments. Across all participants and scanning sessions, fewer than 10% of blocks were excluded due to technical issues (e.g., loss of fixation or tracking instability), and the remaining data showed stable fixation across conditions.

## Author contributions

LL, XCS, and SGK designed the experiments. XCS ran the experiments. XCS, LL, and ZKDS analyzed the data. XCS and LL wrote the first draft. All revised the paper. We declare no competing financial interests.

## Acknowledgements

This study was supported by research grants from the National Natural Science Foundation of China (32161133009, 32071041), Shanghai Science and Technology Committee (20ZR1439500), China Ministry of Education (ECNU 111 Project, Base B1601), and NYU Shanghai (the major grant seed fund and the boost fund).

