## Supplementary Materials for "Distinct neural processing of real versus unreal optic flow in the human brain"

### *Decoding accuracy as a function of voxel numbers*

To illustrate how the mean decoding accuracy, averaged across all non-zero coherence levels, varies with the number of selected voxels, we used the SVM-RFE-CBR algorithm to rank and select the most informative voxels, ranging from 50 to 700 in steps of 10. This analysis was conducted separately for each condition and ROI, providing a detailed understanding of the relationship between decoding performance and voxel number. To examine the differences in decoding accuracy between expansion and contraction of real and unreal optic flow, we conducted separate paired *t*-tests for each ROI, as shown in **Figure S1**.

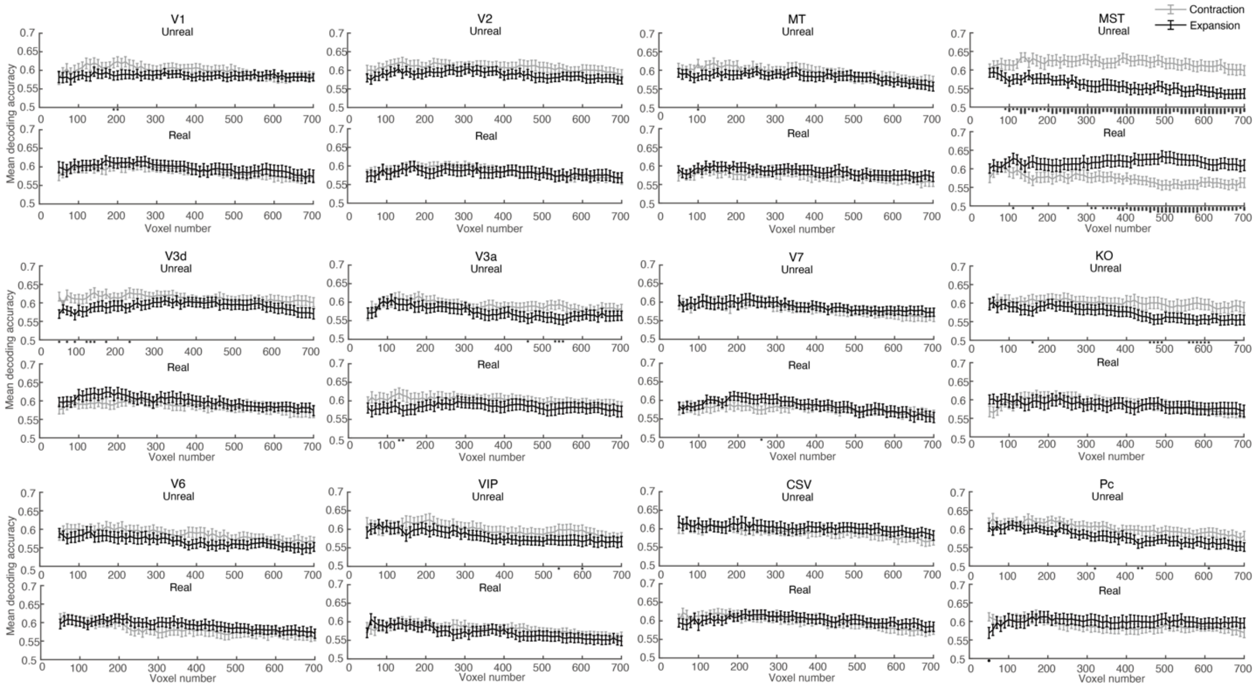


**Figure S1**. Mean decoding accuracy averaged across all non-zero motion coherence levels as a function of voxel numbers (ranging from 50 to 700 in steps of 10) for contraction (gray lines) and expansion (black lines) motion patterns of the unreal (upper panels) and real optic flow (bottom panels) in each ROI. Asterisks indicate significant differences in decoding accuracy between expansion and contraction motion patterns for the real and unreal optic flow stimuli, as revealed by paired *t*-tests (*: 0.01 ⩽ p < 0.05; **: 0.001 ⩽ p < 0.01; ***: p < 0.001).

### *Eye movement data*

During scanning, participants were instructed to fixate on a fixation point that appeared at the beginning of the trial and maintain their fixation on the center of the display throughout the trial. If participants followed our instructions, the pattern of their eye movements should not vary across the motion stimuli. To confirm that different patterns of brain activations were not due to different patterns of eye movements for different motion stimuli, we recorded participants' eye movements throughout the trial, and compared the average horizontal/vertical position, horizontal/vertical RMSE, and amplitude/number of saccades of eye movements across different experimental conditions (**Figure S2**).

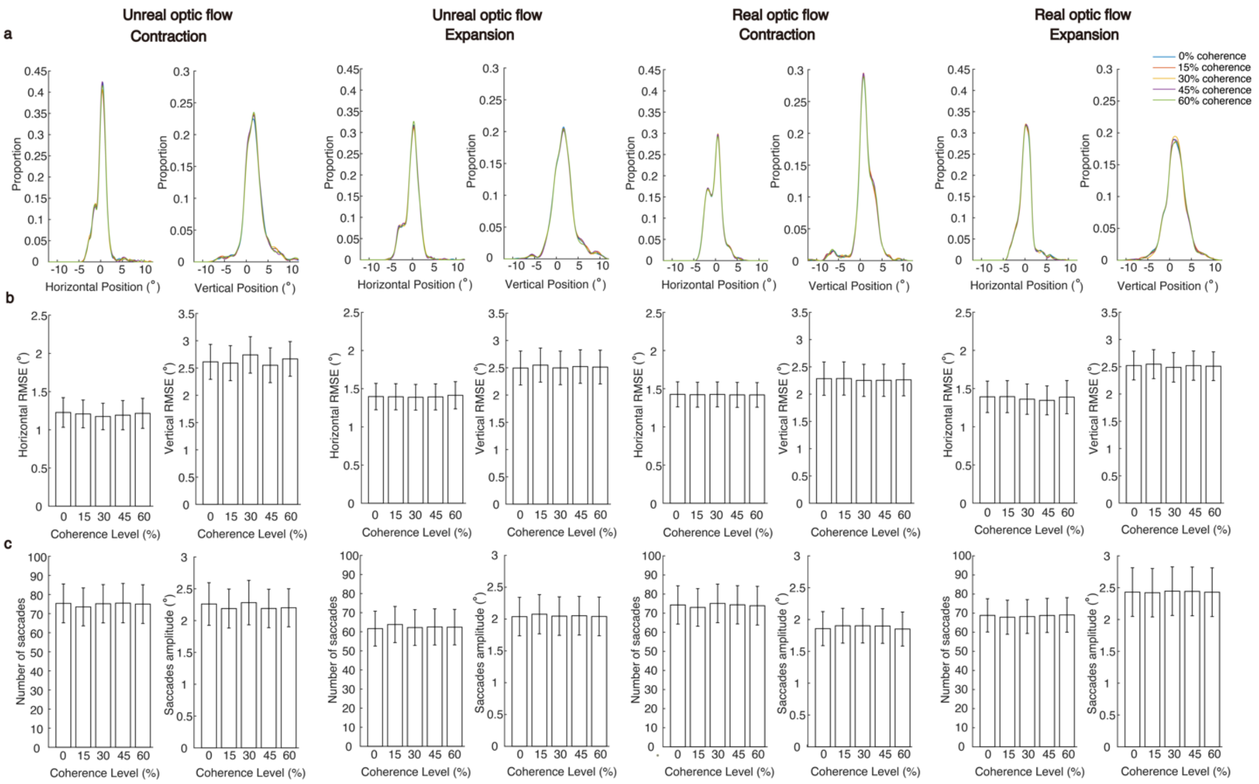


**Figure S2**. Eye movement data during scanning in Experiment 1. **(a)** Proportion of eye position data of 30 participants as a function of the deviation between actual eye fixation and the center of the display along horizontal (each left panel) and vertical (each right panel) direction for the five coherence levels in each experimental condition. The conditions are plotted from left to right: unreal optic flow-contraction, unreal optic flow-expansion, real optic flow-contraction and real optic flow-expansion. **(b)** Horizontal (left) and vertical (right) RMSE against the five coherence levels for each experimental condition. **(c)** Saccade amplitude and the number of saccades against the five coherence levels for each experimental condition. Error bars indicate ±1 SE across 30 participants

**Figure S2a** plots the overall proportion of horizontal or vertical eye positions as a function of the deviation between the actual eye fixation position and the center of the display for each motion coherence level in each experimental condition. Similar trends were observed across different coherence levels in each experimental condition. **Figure S2b** plots the horizontal or vertical RMSE averaged across participants against the five coherence levels for each experimental condition. A one-way repeated-measures ANOVA on horizontal or vertical RMSE was conducted for each experimental condition. The results revealed no significant main effect of coherence level for either horizontal (*p*s > 0.18) or vertical RMSE (*p*s > 0.32) across all experimental conditions. **Figure S2c** plots the amplitude and number of saccades averaged across participants against the five coherence levels for each experimental condition. Likewise, a one-way repeated-measures ANOVA on the amplitude or number of saccades revealed no significant main effect of coherence level was found for either saccade amplitude (*p*s > 0.058) or number of saccades (*p*s > 0.11) across all experimental conditions. These findings suggest that there were no significant differences in patterns of eye movements, such as horizontal/vertical RMSE and amplitude/number of saccades, across different motion coherence levels in each experimental condition.

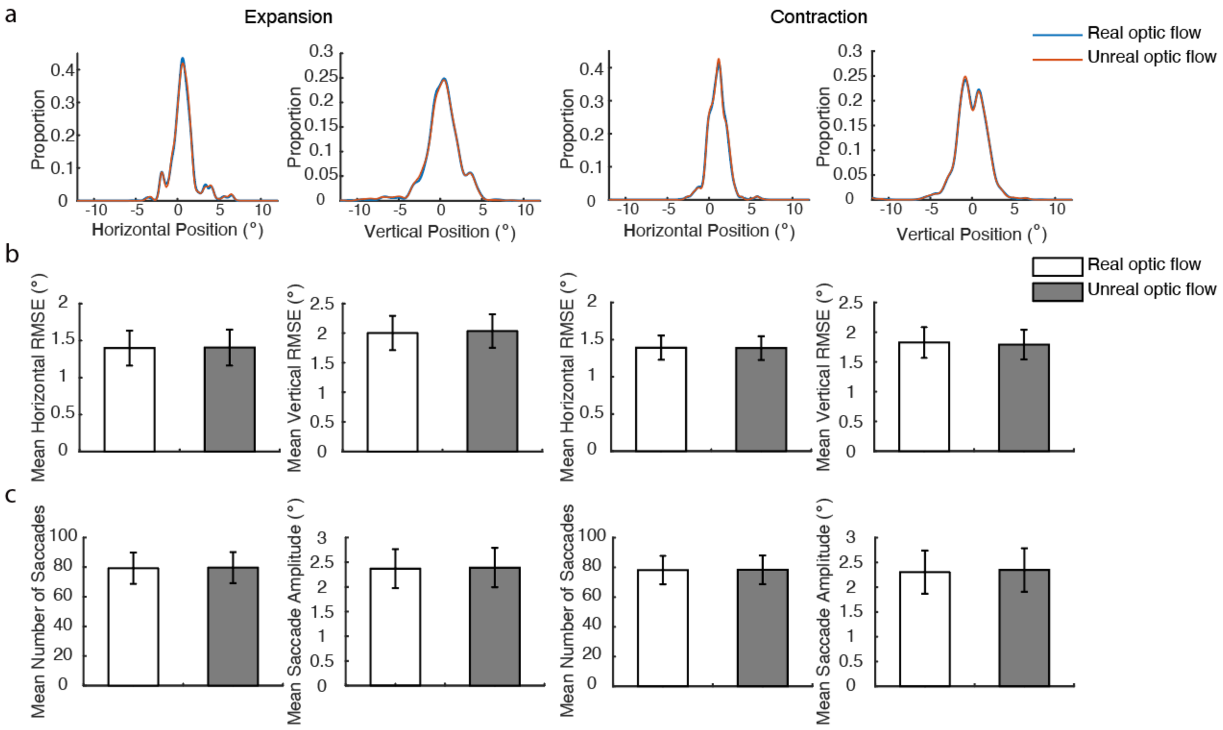


**Figure S3**. Eye movement data during scanning in Experiment 2. **(a)** Proportion of eye position data of 20 participants as a function of the deviation between actual eye fixation and the center of the display along horizontal (each left panel) and vertical (each right panel) direction for the real and unreal optic flow in each motion pattern condition. **(b)** Horizontal (left) and vertical (right) RMSE against the real and unreal optic flow for each motion pattern. **(c)** Saccade amplitude and the number of saccades against the stimulus types for each motion pattern. Error bars indicate ±1 SE across 20 participants

**Figure S3a** shows the overall distribution of fixation positions relative to the center of the display along the horizontal and vertical axes under real and unreal optic flow conditions for both expansion and contraction motion patterns. The distributions of fixation offsets were highly similar across all experimental conditions. **Figure S3b** presents the RMSE of eye position in the horizontal and vertical directions for each condition. Separate 2 (stimulus type: real vs. unreal optic flow) × 2 (motion pattern: expansion vs. contraction) repeated-measures ANOVAs were conducted for horizontal and vertical RMSE. The results revealed no significant main effects of stimulus type or motion pattern for horizontal RMSE (stimulus type: $F(1, 19) = 0.0040$, $p = 0.95$, $\eta^2 = 0.00021$; motion pattern: $F(1, 19) = 0.0040$, $p = 0.95$, $\eta^2 = 0.00021$), nor for vertical RMSE (stimulus type: $F(1, 19) = 0.39$, $p = 0.54$, $\eta^2 = 0.020$; motion pattern: $F(1, 19) = 0.0055$, $p = 0.94$, $\eta^2 = 0.00029$). The interaction between stimulus type and motion pattern was also not significant for either horizontal RMSE ($F(1, 19) = 0.20$, $p = 0.66$, $\eta^2 = 0.010$) or vertical RMSE ($F(1, 19) = 3.20$, $p = 0.090$, $\eta^2 = 0.14$). **Figure S3c** shows the mean number and amplitude of saccades across experimental conditions. Similar 2 × 2 repeated-measures ANOVAs were conducted for both measures. No significant main effects were observed for saccade count (stimulus type: $F(1, 19) = 0.016$, $p = 0.90$, $\eta^2 = 0.00086$; motion pattern: $F(1, 19) = 0.22$, $p = 0.65$, $\eta^2 = 0.011$) or saccade amplitude (stimulus type: $F(1, 19) = 0.011$, $p = 0.92$, $\eta^2 = 0.00056$; motion pattern: $F(1, 19) = 1.99$, $p = 0.17$, $\eta^2 =$

0.095). No significant interaction effects were found for either saccade count ($F(1, 19) = 0.022$, $p = 0.88$, $\eta^2 = 0.0011$) or saccade amplitude ($F(1, 19) = 0.22$, $p = 0.64$, $\eta^2 = 0.012$).

***Localizer scans***

In a separate session of the fMRI experiment, several localizer scans were conducted to define V1/V2/V3d/V3a/V7, MT/MST, KO/V3b, and V6/VIP/CSv/Pc ROIs for each hemisphere of each participant on their reconstructed cortical mesh. V1/V2/V3d/V3a/V7 ROIs were defined by the standard retinotopic mapping procedures with rotating wedge stimuli (see **Figure S4**) (DeYoe et al., 1996; Engel et al., 1994; Sereno et al., 1995; Swisher et al., 2007). MT and MST were defined by the use of an ipsilateral stimulus (see **Figure S5**) based on the method used by Huk et al. (2002). V3b/KO was defined by stimuli with vs. without motion boundary (see **Figure S6**) (Dupont, 1997; Zeki et al., 2003). V6/CSv/VIP/Pc were defined by flow filed with one center of motion (CoM) vs. with nine CoMs (see **Figure S7**) as described in the cited studies (Cardin & Smith, 2010; Wall & Smith, 2008). Further details are provided below.

***Localizers for V1/V2/V3d/V3a/V7***. One functional scan was acquired to define the ROIs for each quadrant of early retinotopic visual areas (V1 and V2), and for the V3d, V3a, and V7 hemifield representations. This set of ROIs was intended to correspond to the underlying visual maps and was defined by the posterior and anterior phase reversals of each hemifield map in the angular dimension (typically anterior/posterior). For all participants, we determined their activation map boundaries based on functional criteria.

The primary mapping stimulus consisted of a static background, viewed through a constantly moving transparent aperture in a dark foreground layer. The background was a static, flickering chromatic radial checkerboard, with the radius of the checkerboard scaled logarithmically to approximate a cortical magnification function (Horton and Hoyt, 1991). Checkerboard luminance alternated spatially between "dark" and "light" checks, in addition to a random chromatic component, to provide both luminance and chromatic contrast. Dark and light checks reversed, and the chromaticity was randomly reassigned, at the 4 Hz flicker frequency (**Figure S4a**). On the polar angle mapping scan, the transparent aperture through which the checkerboard background was viewed comprised a wedge rotating about fixation (**Figure S4b**). Each wedge was 45° in width; thus the checkerboard was divided into 8 polar angle wedges. Each polar angle wedge was presented for 8 s in each circle; thus the periodicity of stimuli was 64 s per circle (12 cycles in total). There was also an 8-s fixation display with no stimulus but a red fixation point (diameter: 1.5°) on the center of a blank screen at the beginning and at the end of the run. The total experiment lasted for 784 s. A red fixation point was presented in the center of the display throughout the task. In 25% of trials, the color of the fixation point was changed to blue. Participants were asked to fixate on the center of the screen and press a button to report the trials containing the blue fixation point.

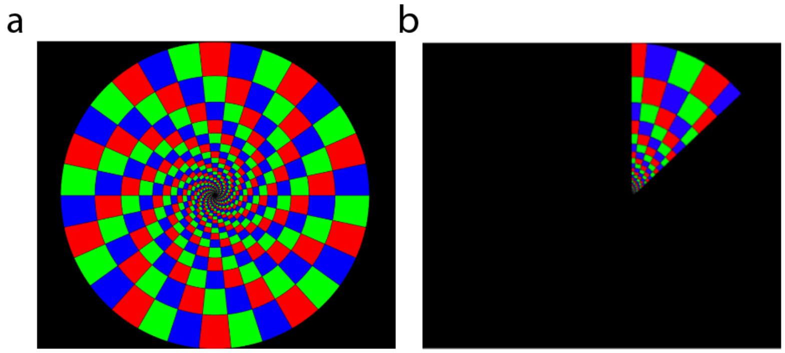


**Figure S4**. Stimuli were used in the retinotopic mapping experiment to localize the V1, V2, V3d, V3a, and V7. **(a)** The static chromatic radial checkerboard sample used in the experiment, with the radius of the checkerboard scaled logarithmically to approximate a cortical magnification function (Horton and Hoyt, 1991). And **(b)** the illustration of a wedge (angle width: 45°) viewed through a constantly moving transparent aperture in a dark foreground layer.

***Localizers for MT/MST***. For all participants, MST was defined (and distinguished from MT) based on the previous studies (Huk et al., 2002). Huk et al. (2002) performed a series of fMRI experiments to divide the human MT+ complex into subregions that may be identified as homologs to a pair of macaque motion-responsive visual areas: the middle temporal area (MT) and the medial superior temporal area (MST). They found that area MT within MT+ did not respond to ipsilateral peripheral stimuli and had a small receptive field. Conversely, area MST responds to peripheral stimuli in both the ipsilateral and contralateral visual hemispheres and has a large receptive field. Thus, the peripheral moving stimuli would be expected to evoke neuronal activity in the contralateral hemisphere in both MT and MST, but in the ipsilateral hemisphere only in MST, where the receptive fields are large enough to extend into the ipsilateral hemifield (e.g., Dukelow et al., 2001; Huk et al., 2002). The central moving stimuli relative to the static stimuli would be expected to evoke neuronal activity in the MT+ complex. With this procedure, MST can be identified in terms of the presence of ipsilateral responses, and MT can be identified based on the remaining regions of the MT+ complex other than MST.

Four types of stimuli were tested: (1) Static high contrast 702 white dots (diameter: 0.08°; luminance contrast: 100%) randomly distributed in a circular field of 19° × 19° (**Figure S5a**). Radial motion patterns in which dots moved (8.6°/s) alternately inwards and outwards along the radial axes, thus creating alternating contraction and expansion. Inwards and outwards motion were alternated every one second. The dots were restricted to a peripheral circular aperture (diameter: 19°) whose center was located either (2) at the center of the screen at the fixation point (**Figure S5b**) or (3) 6.3° to the left (**Figure S5c**) or (4) right of the fixation point (**Figure S5d**). Thus, radial motion stimuli were restricted to the center of the screen or to the left or right hemisphere. This task had 40 blocks (4 conditions × 10 blocks) and 2 fixation blocks. Each stimulus block contained 8 trials of 2-s expansion/contraction motion stimulus. The testing order of stimulus was randomized. Each fixation block had 8 trials of 1-s display with no stimulus but a red fixation point (diameter: 0.2°) on the center of a blank screen at the beginning and at the end of the task. The total experiment lasted for 656 s. A red fixation point was presented in the center of the display throughout the task. In 25% of trials, the color of the fixation point was changed to blue. Participants were asked to fixate on the center of the screen and press a button to report the trials containing the blue fixation point.

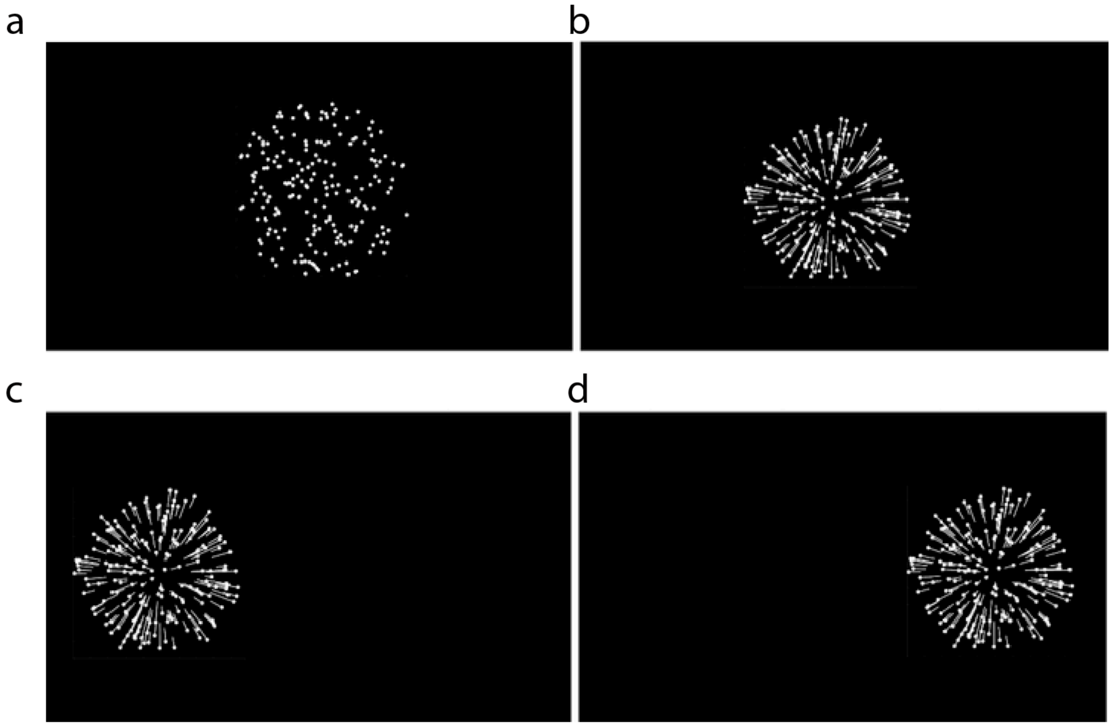


**Figure S5**. Stimuli were used to localize the MT and MST. Four types of stimuli were tested: **(a)** The static white dots whose center was located at the center of the screen. And the radial motion pattern whose center was located **(b)** at the center of the display or **(c)** to the left or **(d)** to the right hemisphere. In **(b)-(d)**, the white dots indicate the dot positions at the end of a trial. The white lines represent the dot motion trajectories in the trial.

***Localizers for V3b/KO***. The kinetic occipital region (KO) was identified in all participants using stimuli similar to those originally used to define KO (Dupont et al., 1997; Zeki et al., 2003). Area KO was shown to process both shape and motion information, the conjunction of which is typically present in kinetic contours. We used two conditions including presentation of kinetic (GKIN) (**Figure S6a**) and uniform motion (UNIFORM) pattern (**Figure S6b**). The kinetic gratings and uniform motion were presented with pixels moving along horizontal, vertical, and both oblique (45° and 135°) axes within the square field (diameter: 5.3°) against the gray background (diameter: 19°). In these stimuli, the pixels moved backward and forward along a single axis for a total of 300 ms before switching randomly to another axis. The rate of change in motion direction was also exactly the same in kinetic gratings and uniform motion stimuli. This task had 16 stimulus blocks (2 conditions × 8 blocks) and 5 fixation blocks. Each stimulus block contained 20 trials of an 800-ms grating stimulus. The testing order of stimulus blocks was randomized. Each fixation block also had 20 trials of an 800-ms display with no stimulus but a white fixation point (diameter: 0.2°) on the center of a blank screen at the beginning, in the middle, and at the end of the run (a fixation block was presented every four stimulus blocks). The total experiment lasted for 336 s. In this task, participants remained passive to the visual stimulus, fixated on the white fixation point, and pressed a button when the color of the fixation point changed to gray in 25% of trials.

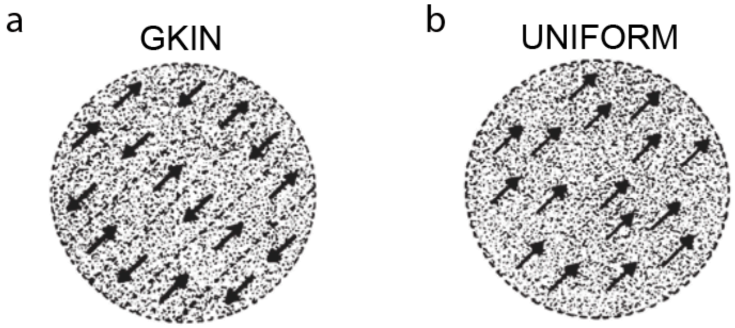


**Figure S6**. Stimuli used to localize the KO, and were exactly the same as those used in the previous study (Dupont et al., 1997). Two types of stimuli were tested: **(a)** kinetic grating generated by pixels of the randomly textured pattern moving in opposite directions, parallel to the boundary. **(b)** The same randomly textured pattern moving uniformly in a single direction. The figure was adapted from the study by Dupont et al., 1997.

***Localizers for V6/VIP/CSv/Pc***. For all participants, the V6, CSv, VIP, and Pc were defined based on the previous studies (Wall & Smith, 2008). These areas have been shown to respond strongly to a single optic flow stimulus (i.e., single, ego-motion-compatible patch) (**Figure S7a**) but become relatively unresponsive when the stimulus is surrounded with further flow patches (i.e., multiple, ego-motion-incompatible patches) (**Figure S7b**) and thereby made inconsistent with ego-motion.

All stimuli consisted of high contrast, moving, random dot patterns on a dark background. Each white dot (diameter: 0.09°; contrast level: 100%) moved in a straight path at a speed of 4.3°/s for a lifetime of 600 ms before disappearing and reappearing at a new, random location. Global patterns of optic flow were produced by control of the local motion directions of the dots. The experiment contained two stimulus conditions: the ego-motion-compatible condition composed of a single flow patch, and the ego-motion-incompatible condition composed of nine flow patches. The ego-motion-compatible condition consisted of a 19° × 19° square field of 585 dots moving in a coherent optic flow pattern containing expansion/contraction and rotation components that varied over time, consistent with self-motion on a varying spiral trajectory. The ego-motion-incompatible condition consisted of the same 19° × 19° stimulus area divided into nine identical panels, each containing a moving stimulus similar to ego-motion-compatible condition but smaller. The dot size, dot speed, and number of dots in the whole array were made identical across conditions. This task had 20 stimulus blocks (single/nine patch conditions × 10 blocks) and 2 fixation blocks. Each stimulus block contained 8 trials of a 2-s expansion/contraction rotation motion stimulus. The testing order of stimulus blocks was randomized. Each fixation block had 16 trials of a 1-s display with no stimulus but a red fixation point on the center of a blank screen at the beginning and at the end of the task. This task lasted 352 s in total. A red fixation point (diameter: 0.2°) was presented in the center of the display throughout the task. In 25% of trials, the color of the fixation point was changed to blue. Participants were asked to fixate on the center of the screen and press a button to report the trials containing the blue fixation point.

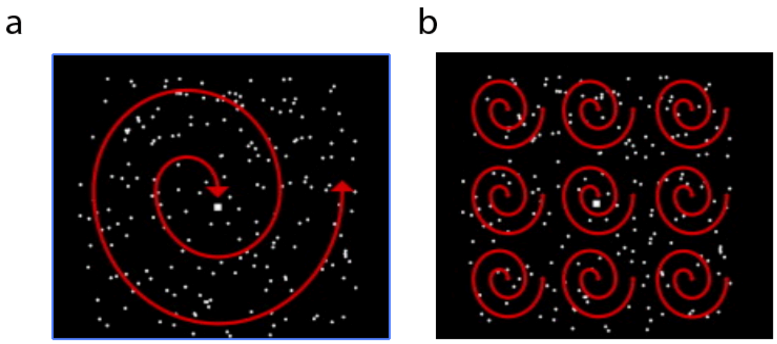


**Figure S7**. Stimuli were used to localize the V6, CSv, VIP, and Pc, and were the same as those used in the previous study (Wall & Smith, 2008). The stimulus was either **(a)** a single patch of time-varying optic flow or **(b)** an array of nine similar patches.

For all participants, the ROIs identified in **Figures S4-7** were subsequently used for the main fMRI analyses in the present study, ensuring that all activation and decoding results were based on individually defined, functionally localized brain regions. The average Montreal Neurological Institute (MNI) coordinates of the ROIs identified in Experiment 1 for the 30 participants are reported as mean ± standard error separately for the left and right hemispheres (see **Supplementary Table 1**). These coordinates represent the average central locations of each ROI across participants and provide a reference for group-level analyses as well as a basis for comparison across studies.

**Supplementary Table 1**. Mean MNI coordinates of functionally localized regions of interest (ROIs).

| ROIs | MNI coordinates | | | | | |
|---|---|---|---|---|---|---|
| | left | | | right | | |
| | x | y | z | x | y | z |
| V3B/KO | -32±4 | -87±4 | 10±5 | 36±4 | -82±4 | 10±5 |
| V6 | -13±3 | -71±5 | 25±6 | 15±3 | -70±5 | 28±6 |
| MT+ | -45±3 | -71±5 | 4±4 | 47±3 | -66±4 | 2±4 |
| MST | -44±3 | -70±5 | 6±5 | 46±4 | -66±5 | 4±5 |
| MT | -45±4 | -71±6 | 2±6 | 47±4 | -66±5 | 0±5 |
| V3a | -21±3 | -88±4 | 21±5 | 26±5 | -84±3 | 22±5 |
| CSV | -8±2 | -14±7 | 46±5 | 9±2 | -17±7 | 47±5 |
| Pc | -11±2 | -39±6 | 48±6 | 11±2 | -41±6 | 51±5 |
| VIP | -31±5 | -53±8 | 51±6 | 31±5 | -51±8 | 52±4 |
